# DMEM, or Opti-MEM, that is the Question: An Important Consideration for Extracellular Vesicle Isolation and their Downstream Applications

**DOI:** 10.1101/2025.11.17.683537

**Authors:** Nikki Salmond, Jacob Melamed, Sina Halvaei, Karla C. Williams

## Abstract

Serum-free synthetic media are frequently used as an alternative to extracellular vesicle-depleted serum containing media (EV-DEP) for EV production. However, the impact of this medium on EV biogenesis, release, and composition remains poorly understood. Here, we comprehensively characterised EV release by MDA-MB-231 or HEK-293T cells cultured in EV-DEP DMEM versus a serum-free synthetic medium, Opti-MEM. In Opti-MEM culture, cells released significantly more CD9- and CD63-positive EVs compared to EV-DEP DMEM. Proteomic analysis revealed that EVs from EV-DEP DMEM contained more histones and bovine proteins. Cells cultured in Opti-MEM released a substantially higher proportion of small-EVs derived from the sphingomyelinase-MVB pathway (60%) compared to EV-DEP DMEM (30%). Conversely, cells cultured in EV-DEP DMEM were more reliant upon the ROCK kinase pathway to release ectosomes (45 %) compared to OptiMEM (15 %). Where cells cultured in EV-DEP DMEM employed RabGTPase-dependent mechanisms for MVB-derived EV release (Rab3d, Rab27a, and Rab27b), cells cultured in Opti-MEM did not. Given that cells cultured in Opti-MEM vs EV-DEP DMEM produce EVs with different protein signatures from distinct molecular pathways, the choice of medium should be carefully considered when designing EV studies – particularly if they are to be used in therapeutic or immunological experiments.

## Introduction

Extracellular vesicles (EVs) are membrane-limited biological nanoparticles released by all cell types into the culture media, under standard culture conditions, and into the extracellular environment and biological fluids in vivo^1^. Cell lines and primary cells can be cultured in 2D on plastic plates, in 3D spheroids and organoids, in suspension, or in large bioreactors. Regardless of the technique by which cells are grown, cell culture media are required to provide the cells with growth factors, amino acids, vitamins and sugars needed for cellular growth^2^. EVs are harvested by laboratories worldwide at small-scale and large-scale for various applications including basic scientific research to investigate cell communication, immunology, pathology, and health, as well as in clinical research for biomarker discovery and therapeutic development^3,4^.

The EV field is continuously and rapidly evolving in several sectors including, basic science research, biomarker discovery, therapeutic delivery, and, importantly, methodology development to improve strategies for the production and isolation of EVs. EV production and isolation experiments are multi-step processes that demand significant time commitment^5^. Preparing and executing EV collections often requires intensive cell culture procedures including the plating of large numbers of cells, with a significant expenditure of cell culture plastic ware and media, all of which becomes more intensive as the scale of EV production and isolation increases. Large-scale EV production protocols are used by many researchers and are particularly important in the development of EV-based therapeutics where high yield and purity with low hands-on time for EV preparation and isolation is largely desired^4^. Any method development that can increase EV yield and purity while decreasing the experimental hands-on time would represent a welcomed advancement for the EV community.

A further consideration for the production of EVs, is the purity at which they are isolated. A major consideration for achieving EV preparations of reasonable purity is not just the choice of EV isolation technique, but also the cell culture media used for cell maintenance and growth. Traditionally, researchers use fetal bovine serum to supplement cell culture media as the growth factors, lipids, and amino acids present in the serum support cell growth and health^2,6,7^. For use in EV experiments, fetal bovine serum needs to be depleted of bovine-EVs to prevent contamination of cell line-derived EVs of interest. Preparation of EV-DEP serum requires an overnight ultracentrifugation spin or a filtration process, increasing the equipment needs, length of experiment, and the hands-on experimental time necessary to prepare for an EV collection experiment^8^. While EV-depletion decreases the number of fetal bovine EVs in the media, it does not completely remove all of them, thereby, some residual contamination may still occur^9^. Furthermore, EV depletion of FBS depletes other factors present in serum important for cell growth and physiological phenotypes – many researchers have reported impaired proliferation^10^, viability^9,10^, migration^11^, and immune cell responses^12,13^ in media containing EV-depleted (EV-DEP) serum^8,9^.

Further complicating EV studies, fetal bovine serum proteins present in EV-depleted media can form protein coronas on cell line-derived EVs which may impact EV biology and function. Fetal bovine serum also contains lipoproteins, protein aggregates, RNA, and DNA, all of which co-isolate with EV preparations, are difficult to completely remove, and change EV uptake/functionality^6,8,12,14–16^. This can be a serious issue depending on the downstream applications of EVs and is undesirable for use of the EVs in immune assays or for therapeutic development. To eradicate the possibility of bovine contaminates co-isolating with EVs of interest, the use of serum-free and synthetic media is becoming a more common choice. In recent years, there have been improvements in the production and cost of synthetic media improving laboratory access to these previously limited and expensive products. Researchers are now commonly using serum-free synthetic media as an alternative to EV-DEP serum supplemented media for EV collections.

The selection of cell culture media, while important, may not always be clear, and selection is often a user-dependent decision with consideration of the final EV product and their downstream applications. Here, we aimed to further our understanding on the impact of different mediums (10% EV-DEP serum-supplemented DMEM versus serum-free Opti-MEM) on EV release and EV characteristics from commonly used cell lines (HEK-293T and MDA-MB-231). Our findings highlight that cells cultured in Opti-MEM produce a higher yield of sphingomyelinase-MVB derived EVs with minimal histone contamination and no bovine protein contamination compared to EV-DEP serum DMEM cell culture conditions. Cells cultured in EV-DEP DMEM produced a lower yield of EVs, primarily originating from ROCK kinase dependent ectosome biogenesis routes, as well as sphingomyelinase and RabGTPase MVB-dependent pathways. However, these EV preparations were rich in histone and bovine proteins which may be undesirable contaminates for therapeutic applications. Our study provides a framework to guide an informed decision when selecting a cell culture media for EV applications.

## Materials and Methods

### Cell lines and cell line generation

HEK293-T and MDA-MB-231 cells were purchased from ATCC. The myr-palm-mNeonGreen(2x) vector was a gift from Salvatore Chiantia (Addgene plasmid # 197919)^17^ and was used to transfect HEK-293T cells using lipofectamine 3000 according to manufacturer’s instructions (Invitrogen) to generate stably transfected HEK-293T-myr-palm-mNeonGreen(2x) cells maintained using antibiotic selection in 1 mg/mL Neomycin (G-418) (Cayman Chemical). The pLC-ZsGreen-P2A-Hygro vector was a gift from Eva Gottwein (Addgene plasmid # 124301)^18^ and was used to transfect MDA–MB-231 cells using lipofectamine 3000 according to the manufacturers protocol to generate stably transfected MDA–MB-231–ZsGreen cell lines maintained by selection in 500 µg/mL Hygromycin B (PlantMedia). Stably transfected cells were trypsinised, washed with PBS twice and run on the flow cytometer (CytoFLEX-S, Beckman Coulter) to confirm their successful transfection. Cell lines that showed less than 90% FITC positive population, were sorted for the top 20% highest expressors using the Astrios cell sorter (Beckman Coulter).

### Lentivirus production and transduction

The pLenti-PalmGRET plasmid was a gift from Charles. P Lai (Addgene plasmid #158221)^19^ and was used to produce lentivirus particles for transduction of an MDA-MB-231-PalmGRET stable cell line maintained in 4 µg/ml Puromycin. Lenti-X™ HEK-293T cells (Takara Bio) were plated in a 6 well plate in 10 % heat inactivated FBS DMEM (HI-DMEM) to achieve 70 % confluence the following day when they were transfected using lipofectamine 3000 according to manufacturer’s instructions with 900 ng psPAX2, 100 ng pMD2.G, and 1 µg pLenti-PalmGRET^19^ for lentivirus production. Twenty-four hours later, media on transfected LentiX-HEK-293 cells was refreshed with 2.5 ml HI-DMEM and lentivirus producing cells conditioned media for a further 24 hours. Conditioned media was harvested, spun at 300 x g for 10 minutes, then 2500 x g for 15 minutes to remove floating cells and debris. Clarified media containing PalmGRET-lentivirus was then used to transduce MDA-MB-231 cells. For transduction, 4,800,000 MDA-MB-231 cells were mixed with 13.2 µl polybrene and 1.5 ml lentivirus conditioned media in a 10 cm plate in a total volume of 12 ml HI DMEM. After 24 hours media was refreshed, and 24 hours later cells were put under puromycin selection 4 µg/ml. Surviving cells stably expressing PalmGRET were bulked up and stocks frozen down.

### Cell culture

All cells were cultured and maintained between 20 – 80% confluence at 37 °C, 5% CO_2_ in 10% FBS (Gibco) DMEM (Wisent). For EV collections or nanoflow analysis, cells were maintained in either 10 % FBS DMEM, 10% EV-DEP serum-supplemented DMEM, or OptiMEM-GlutaMax (Thermo Fisher Scientific).

### Transient expression of H2B-zsGreen

The ZsGreen1-H2B-6 vector was a gift from Michael Davidson (Addgene plasmid # 56514) and used to transiently transfect HEK-293T cells in a 6-well plate using lipofectamine 3000 according to the manufacturers protocol. Twenty-four hours later, cells were re-plated for the analysis of EV release.

### Flow Cytometry of cells

MDA-MB-231-zsGreen cells were lifted by trypsinisation and washed twice in PBS before analysis of ZsGreen expression by flow cytometry (CytoFLEX-S, Beckman Coulter) using default forward side scatter settings and FITC gain 10 for detection of ZsGreen positive cells. The zsGreen fluorescent profile of cells incubated in Opti-MEM versus EV-DEP DMEM was compared on a histogram.

### Plating of cells for EV release quantitation – 96 well plates

MDA-MB-231-zsGreen and Hek-293T-myr-palm-mNeonGreen(2x) cells were trypsinised, quenched in DMEM and washed twice in PBS before being plated in triplicate in 96 well plates in the appropriate media or treatment conditions. During optimization, 5,000, 10,000, 20,000, or 40,000 cells per well were plated in either Opti-MEM or 10% serum-supplemented DMEM. After optimization, for histone2B-zsGreen overexpression experiments, 20,000 cells per well were plated per well in Opti-MEM or 10% EV-DEP serum DMEM. After plating, cells were allowed to condition media for 48 hours (optimization experiments) or 24 hours (Histone2B-zsGreen experiments) before determining cell confluence (IncuCyte SX5) and analysing the number of EVs in 50 µl conditioned media diluted in 50 µl 0.02 µm filtered PBS using nanoflow cytometry (CytoFLEX-S, Beckman Coulter), the protocol is detailed in the ‘Nanoflow cytometry’ section below.

### Plating of cells for small-molecule manipulation of EV release – 12 well plates

For treatment of cells with inhibitors and stimulators of EV release, 150,000 MDA-MB-231-PalmGRET cells were plated in each well of a 12 well plate in DMEM 10% FBS. Twenty-four hours later, cells were washed 2 times with PBS and media was replaced with 1.5 ml Opti-MEM or EV-DEP DMEM containing bafilomycin (160 nm), GW4869 (5 µM), Y-27632 (5 µm), or DMSO matched control (1:1000). The next day, cell confluence was quantified by imaging using IncuCyte SX5 and conditioned media was harvested and clarified using a 300 x g spin for 10 minutes, and a 2,500 x g spin for 15 minutes. A final 10,000 x g spin was done to pellet large EVs which were re-suspended in 200 µl PBS. The remaining conditioned media was then used to analyse the remaining small EVs. Large EVs were analysed after diluting 10 µl in 90 µl PBS, and small EVs were analysed using 50 µl conditioned media in 50 µl PBS using the CytoFLEX protocol detailed in the ‘Nanoflow cytometry’ section.

### Plating of cells for siRNA knockdown of RabGTPases – 12 well plates

Three hundred thousand MDA-MB-231-PalmGRET cells were plated in each well of a 12 well plate, the next day cells were transfected with siRNA (25 nM final concentration) to Rab3d, 7a, 11b, 27a&b, and 35 using Lipofectamine 3000 according to manufacturer’s instructions. Twenty-four hours later, cells were trypsinised and replated into two individual 12 wells and allowed to recover for 24 hours at which point cells were washed 2 times with PBS before replenishing with either 1.5 ml Opti-MEM or EV-DEP DMEM. Cells were allowed to condition media between 48-72 hours post-transfection, before cell confluence analysis, and conditioned media clarification as described in ‘plating of cells for small molecule manipulation of EV release’ above. EV quantification is described further in the ‘Nanoflow cytometry’ section.

### Nanoflow cytometry

Samples were set up in a 96 well plate with a PBS wash between each sample. Samples were run slowly for 40 seconds, triggered using 405 nm violet side scatter (1027 detection threshold, gain 100) and the number of ZsGreen, myr-palm-mNeonGreen(2x), or PalmGRET positive EVs were quantified using the FITC setting at a gain of 500. Manual gating was applied to the population of interest and time of interest (10-40 seconds to avoid the first 10 seconds where flow rate can take time to stabilise). The number of events detected within the gate was taken as a representative number of EVs quantified in each different treatment condition and was divided by the cell confluence to account for any changes in cell proliferation.

### EV lysis nanoflow cytometry

To assess the lysis of EVs from H2B-zsGreen expressing HEK-293T and PalmGRET expressing MDA-MB-231 cells, EV or conditioned media was diluted 1:1 in 1 % Triton-x for 10 - 30 minutes before analysing lysis efficiency using nanoflow cytometry.

### IncuCyte analysis of confluence

The confluence of cells was determined by IncuCyte SX5 (Sartorius) imaging of each well. Four images per well were taken per 96 well, or 49 images per well for 12 well plates, and the average confluence was used to determine the number of EVs released/1% cell confluence.

### Cell count and viability

Cells were counted and viability assessed using a Luna II automated cell counter and trypan blue according to manufacturer’s instructions (Logos Biosystems).

### EV-DEP serum preparation

EV-DEP serum was prepared by ultracentrifugation of 20% serum DMEM for 16-18 hours at 100,000 x g, 4 °C. The resultant supernatant was harvested, 0.2 µm sterile filtered and diluted to 10% using serum-free DMEM.

### EV collections

For EV collections, cells were trypsinised and 20 million cells per condition were washed twice with PBS by centrifugation at 300 x g. The cells were then resuspended in 100 mL of 10% EV-DEP serum-supplemented DMEM or Opti-MEM and plated in a Nunc TripleFlask (Thermo Fisher Scientific) for 48 hours. Upon EV harvest, conditioned media was centrifuged at 300 x g for 10 minutes. As HEK-293T cells became less adherent in Opti-MEM, the number of cells from which EVs were isolated were counted in the conditioned media 300 x g spin pellet, and the cell wash supernatant as well as the trypsinised cells. The conditioned media was then further cleaned up by a 2500 x g spin for 15 minutes. Large EVs were removed using a 10,000 x g spin for 30-minutes at 4 °C, and small-EVs were pelleted at 100,000 x g for 70 minutes at 4 °C. The small EVs were pooled and washed in PBS in a 1.5 mL Eppendorf, before further centrifugation at 100,000 x g for 70 minutes at 4 °C using Eppendorf tube floatation. EVs were resuspended in approx. 30 µL PBS for further downstream analysis.

### CFSE staining of EVs

Once EVs were pelleted using the first 100,000 x g spin, EVs were resuspended in 1 mL PBS and incubated with CFSE (10 mM) (1:2000) at 37 °C for 10 minutes before analysing the number of EVs using nanoscale flow cytometry (diluted 1:1000 PBS). Diluted CFSE was run on the CytoFLEX-S to ensure no background signal was detected.

### Lysate preparation

Cells from each cell culture condition were trypsinised and harvested for lysis in RIPA buffer (Thermo) with HALT protease inhibitor (1:100) (Thermo) on ice for 10 minutes. Lysates were centrifuged at 17,968 x g for 10 minutes at 4° before supernatant was harvested for protein quantitation and western blot.

### Protein quantitation

Cell lysate protein concentration was quantified using Rapid Gold BCA assay (Pierce, Thermo Scientific) according to manufacturer’s instructions. The EV protein concentration quantification was achieved by using 5 µL of an EV preparation in the micro-BCA assay (Pierce, Thermo Scientific) according to manufacturer’s instructions.

### Nanoparticle tracking analysis

The presence of particles in an EV preparation was confirmed by nanoparticle tracking analysis (NTA). Briefly, EVs were diluted 1:1000 and detected on the NanoSight LM10 using a 488 nm laser (Malvern Panalytical). A syringe pump at speed setting 40 was used to achieve continuous flow of EVs through the chamber and three 30 second measurements were acquired. The camera level was adjusted according to sample needs and a detection threshold of 5 was used for analysis. NTA software version 3.3.301 was used.

### DNA quantitation

The DNA concentration in EV preparations was ascertained by using 2 µl of EV preparation on the nanodrop which measured the samples at 260 nm. Instrument was used according to manufacturer’s instructions.

### Zeta potential

The charge of EVs was measured by diluting EVs 1:1000 in PBS and measuring the Zeta potential on Malvern ZetaSizer (Malvern Panalytical) according to manufacturer’s instructions.

### Electron Microscopy

Harvested EVs were fixed in 2% electron microscopy grade PFA. Copper 400 mesh carbon type-B grids were glow discharged at 60 µA for 10 seconds. Five microlitre droplets of samples were applied to glow discharged grids and allowed to sit for 30 secs to 10 mins, depending on sample concentration, before blotting excess with Whatman filter paper. Grids were then negatively stained with 2% uranyl acetate for 1 minute. Samples were imaged on a FEI Tekna Spirit 120kV TEM operated at 80kV.

### Western Blot

The appropriate amount of cell lysate or EVs were diluted in 4 x Bolt LDS sample buffer (Thermo) and 10 x Bolt sample reducing agent (Thermo) (unless CD63 or CD9 were detected, in which case, non-reducing conditions were used). Samples were boiled at 95 °C for 10 minutes before loading onto a 10 or 15 well bis-tris NuPAGE gel (Thermo Fisher Scientific). Proteins were separated by running the gel at 220 V for 30 minutes in MES running buffer (50 mM MES, 0.1% SDS, 50 mM Tris Base, 1 mM EDTA) and using Kaleidoscope molecular weight marker (BioRad). Proteins were transferred onto a 0.45 µm nitrocellulose membrane (Thermo Fisher Scientific) in transfer buffer (190 mM glycine, 25 mM Tris Base, 10% MeOH) for 90 minutes at 30 V. Membranes were blocked in 5% milk in TBS-T (20 mM Tris base, 160 mM NaCl, 0.1% Tween) and incubated with primary antibodies diluted in TBS-T overnight at 4°C with constant agitation. Membranes were washed by three 10-minute washes in TBS-T before incubating with secondary LiCOR antibody IRdye® 680RD or 800CW (1:10,000 in TBS-T containing 1% milk) for 30 minutes room temperature in the dark. Three more 10-minute TBS-T washes removed any residual antibody before imaging using the Sapphire Biomolecular Imager (Azure Biosystems). Images were imported into Image Studio Lite 5.2 for image preparation and densitometry analysis. Antibodies used: Vinculin 1:5000 (Abcam: ab91459). CD63 1:500 (BD Biosciences: 556019). CD9 1:500 (Millipore: CBL162). Calnexin 1:5000 (Abcam: ab22595). Histone2B 1:1000 (Abcam: Ab1790).

### Mass spectrometry of EVs

Four microliters of each lysed sample was reduced and alkylated using 50mM TCEP/ 200mM CAA mixture (Tris(2-carboxyethyl)phosphine hydrochloride and 2-chloroacetamide)^20^ for a final concentration of 5 mM TCEP/ 20mM CAA, diluted with 50 mM ammonium bicarbonate (pH ∼ 8) to correct for pH of the samples, and incubated at 37 °C for 20 min. The resultant solution of proteins was cleaned up and digested with 80 ng of Trypsin (Promega) on beads overnight at 37 °C (1:1 Cytiva Sera-Mag, 80% EtOH binding) using the SP3 workflow as previously described.^21^ Finally, quenching of trypsin was achieved by acidification to pH < 2.5.

The liquid chromatography mass spectrometry analysis was done on a Bruker TIMSTof Pro2, connected to a Nanoelute UPLC with a 25cm Ionopticks Aurora column. Standard method parameters were used and are available in the uploaded archived data (ProteomeXchange identifier: PXD068319). Data was analyzed using MS1 LFQ quantification in MaxQuant (v2.5.2.0), and bioinformatic analysis was performed using the Mass Dynamics (https://massdynamics.com/) web tools.^22^

### Statistics and Software

All data handling was done in excel and graph construction and statistical analysis in GraphPad prism 10.4.1. Data was assessed for normal distribution using a Shapiro-Wilk test. If data was normally distributed, an ordinary one-way ANOVA or unpaired T-test was used. If data was not normally distributed, a Kruskal Wallis ANOVA or Mann-Whitney nonparametric T-test was used.

If data was normalized to control condition, either a one-sample t-test (parametric) or Wilcoxon signed rank test (non-parametric) was used with the hypothetical value inserted as 1. \**p* ≤ 0.05. \*\**p* ≤ 0.01. \*\*\**p* ≤ 0.001. \*\*\*\**p* ≤ 0.0001. Image Studio Lite was used for western blot densitometry analysis.

## Results

### Cells cultured in Opti-MEM release more extracellular vesicles than those in serum-supplemented DMEM

Cell culture media composition can impact EV release dynamics. To determine if the use of a serum-free medium - Opti-MEM - as an alternative to 10% serum-supplemented DMEM affected the dynamics of EV release, the number of EVs released into conditioned media by MDA-MB-231-ZsGreen (Figure 1A & B) and HEK-293T-myr-palm-mNeonGreen(2x) (Figure 1C & D) cell lines was quantified by nanoflow cytometry (CytoFLEX-S, Beckman Coulter). The number of EV counts was normalized to estimate EV release based on confluence and ascertain the number of EVs released per 1% cell confluence. MDA-MB-231-ZsGreen cells released significantly more EVs in Opti-MEM as compared to DMEM at all confluences (Figure 1A & E). HEK-293T-myr-palm-mNeonGreen(2x) cells also released more EVs in Opti-MEM, but this was only significant for the 10,000 and 40,000 cells/well seeding density (Figure 1C & E). The different media did not significantly change the cell confluence of MDA-MB-231-ZsGreen cells except when plated at the most confluent cell number, 40,000 cells/well, where cells grew significantly faster in DMEM (Figure 1B). On the other hand, HEK-293T-myr-palm-mNeonGreen(2x) cells grew significantly faster in DMEM at all confluences except the lowest when cells were plated at 5,000 cells/well (Figure 1D). The different cell culture media conditions altered cell morphology of both cell lines, with cells in Opti-MEM presenting a less well-spread acicular-like morphology (Supplementary figure 1).

**Figure 1:**
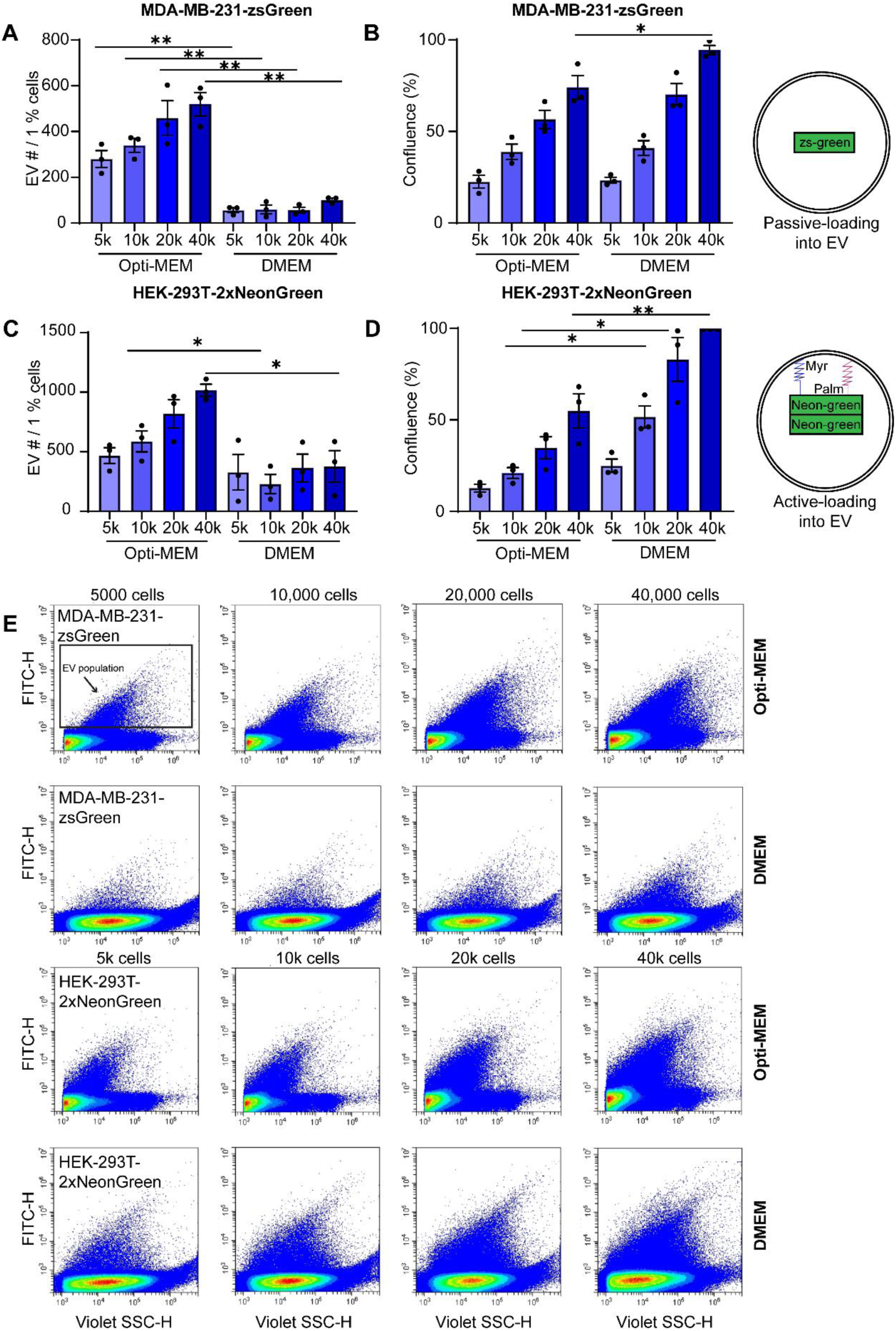
MDA-MB-231-ZsGreen cells and HEK-293T-myr-palm-mNeonGreen(2x) cells release more EVs when cultured in Opti-MEM, compared to 10% serum-supplemented DMEM. 5,000, 10,000, 20,000 or 40,000 MDA-MB-231-ZsGreen (A & B) or HEK-293T-myr-palm-mNeonGreen(2x) (C & D) cells were plated per well, in triplicate, in a 96 well plate in Opti-MEM or 10% FBS-supplemented DMEM. Forty-eight hours later, cell confluence (B & D) was measured by IncuCyte imaging and the number of EVs released (A & C) into 50 µL conditioned media diluted 1:1 in 0.02 µm-filtered PBS, analysed by nanoflow cytometry using CytoFLEX-S. EV number was normalised to cell confluence to take changes in cell proliferation into consideration. Representative EV populations of each cell confluence and media conditioned is shown in (E). N=3. ± SEM. Unpaired T-test.

Representative flow plots of EV populations in each condition are shown in Figure 1E. The EV populations under different cell culture conditions are unique; cells cultured in Opti-MEM produced a compact EV distribution and population shift towards the left, whereas cells cultured in DMEM released a more diffuse population of EVs spread across the center of the flow plot. HEK-293T-myr-palm-mNeonGreen(2x) cells exhibited increased variation in the number of EVs released between experiments likely due to decreased adherence in Opti-MEM, therefore we focused our study on the MDA-MB-231 cells, with supporting data obtained from HEK-293 cells presented in the supplementary data. As EV production can be impacted by multiple variables such as cell seeding density and passage number^23,24^, to minimize noise from inter-experiment variation, data was normalized to control Opti-MEM condition for subsequent nanoflow experiments. To ensure that the increased EV release observed in cells cultured in Opti-MEM was due to increased EV production and not increased expression of ZsGreen in MDA-MB-231 cells, ZsGreen expression was evaluated by flow cytometry and determined to be equivocal in cells cultured in Opti-MEM and 10% serum supplemented DMEM (Supplementary figure 2).

To further characterise the EVs released by MDA-MB-231 cells, EVs were collected from cells cultured in either Opti-MEM or 10% EV-DEP DMEM and analyzed for protein content, charge, nucleic acid content, size distribution, and morphology. The total number of MDA-MB-231 adherent and non-adherent cells at the time of EV collection were not found to be significantly different between culture conditions (Figure 2A, supplementary figure 3B). The total protein concentration of EVs collected from Opti-MEM cell culture conditions was significantly higher than that of EV-DEP DMEM (Figure 2B). Given Opti-MEM culture conditions significantly increase EV number (Figure 1), the increased protein concentration likely reflects the higher EV yield. An evaluation of the approximate protein content per EV identified that the amount of protein per EV trended towards a lower amount for Opti-MEM derived EVs compared to EV-DEP DMEM EVs, although this was not found to be significant (p=0.07) (Figure 2C). To assess the impact of cell culture media on EV charge, the zeta potential of EVs was measured. Cells cultured in Opti-MEM produced EVs that were more negatively charged compared to EV-DEP DMEM (Figure 2D), although this was not found to be significant (p=0.08). To assess changes to nucleic acid content, the DNA content of EV preparations was evaluated and found to be significantly higher in EVs derived from cells cultured in Opti-MEM (Figure 2E). Nanoparticle tracking analysis showed that MDA-MB-231 cells cultured in Opti-MEM released more EVs than those in EV-DEP DMEM (Figure 2F) but the size distribution of the EV populations did not change (Figure 2G). To analyse the number of isolated EVs released by nanoflow cytometry, EVs were stained with CFSE (1:2000). Similar to the results obtained from the MDA-MB-231-ZsGreen cell line (Figure 1), MDA-MB-231 cells cultured in Opti-MEM were found to release significantly more EVs than cells cultured in EV-DEP DMEM (Supplementary figure 4). Electron microscopy showed the presence of EV structures in the micrographs (Figure 2H).

**Figure 2:**
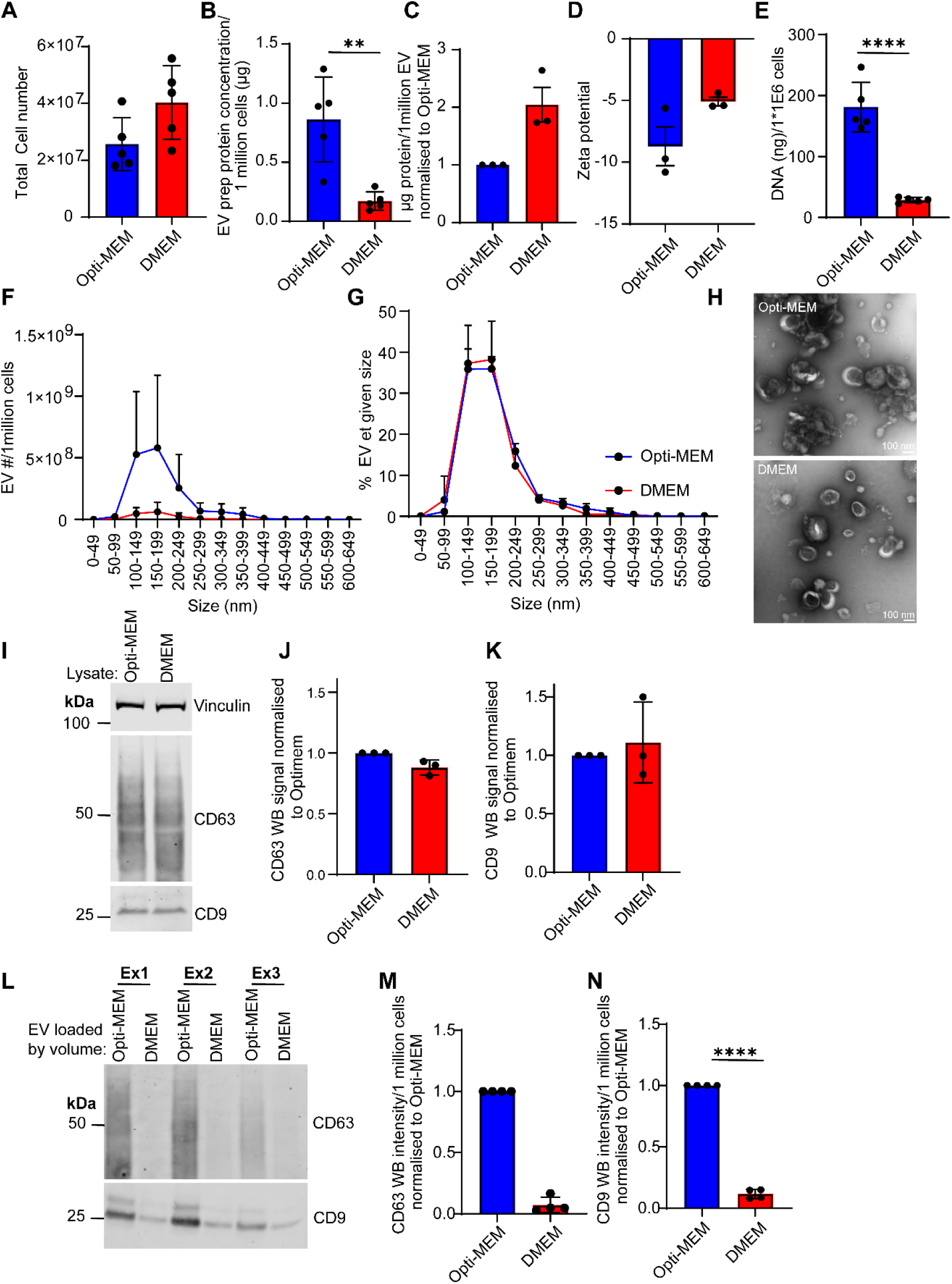
Isolated EVs from MDA-MB-231 cells show that more CD9 and CD63 positive EVs are released by cells when cultured in Opti-MEM media rather than EV-DEP DMEM media. Twenty million MDA-MB-231 cells were plated in the appropriate cell culture media for 48 hours before EV collection by ultracentrifugation. The total cell number (adherent and non-adherent) from which EVs were isolated was recorded N=3. ± SEM (A) and the protein concentration of EV preparation/1 million cells calculated by micro-BCA assay. N=3. *±* SEM. Unpaired T-test. (B). Nanoparticle tracking analysis data was used to calculate the protein concentration/1 million EV N=3. ± SEM. (C). The Zeta potential of EVs diluted 1:1000 in PBS was analysed using a Zetasizer (Malvern) N=3. ± SEM, (D) and DNA concentration/1 million cells recorded using a NanoDrop. N=5. ± SEM. Unpaired T-test. (E). Nanoparticle tracking analysis of EV number/1 million cells (F) and their size distribution (G). N=3. ± SEM. Electron microscopy was used to visualize EVs. N=3. (H). Western blot of 20 µg MDA-MB-231 cell lysate to analyse expression of CD63 and CD9 in Opti-MEM and EV-DEP DMEM (I) with densitometry analysis (J & K). Two microliters of each EV preparation was loaded for western blot analysis of CD9 and CD63 content (L) and densitometry analysis allowed determination of the amount of CD63 and CD9 in EVs released per 1 million cells N=4. + SEM. One sample T-test (CD9) or one sample Wilcoxon test (CD63) (M & N).

Next, EV marker, CD9 and CD63, expression was assessed by western blotting of lysates from MDA-MB-231 cells cultured in Opti-MEM and EV-DEP DMEM. Equal protein concentration of lysates were loaded (20 µg) and no significant difference in the expression of CD9 or CD63 was found indicating that, at the cellular level, the protein abundance of these canonical EV markers was similar in each culture condition (Figure 2 I, J & K). EVs isolated from MDA-MB-231 cells were then loaded by equal volumes, and densitometry results were corrected to cell number to represent the amount of CD9 vs CD63 positive EVs released per 1 million cells. MDA-MB-231 cells cultured in Opti-MEM released more CD9 and CD63 positive EVs per 1 million cells as compared to EV-DEP DMEM (Figure 2 L, M & N), further highlighting the increased EV production in Opti-MEM culture conditions.

To extend our analysis to additional cell lines, EVs isolated from HEK-293T cells in Opti-MEM or EV-DEP DMEM were characterised. HEK-293T cells grew significantly faster in DMEM (Supplementary figure 3A), and, when cultured in Opti-MEM, a greater proportion of the cells were non-adherent compared to the MDA-MB-231 cell line (Supplementary Figure 3B). Both HEK-293T and MDA-MB-231 cells cultured in Opti-MEM or EV-DEP DMEM were assessed for viability and no significant differences were found for adherent cells (Supplementary Figure 3C), but non-adherent MDA-MB-231 cells were found to have a lower viability in DMEM compared to Opti-MEM (Supplementary Figure 3 D & E). The altered adhesion properties of HEK-293T cells in Opti-MEM may lead to inaccurate cell counts and could account for the larger inter-experiment variability observed relative to MDA-MB-231 cells. The protein concentration of EV preparations from HEK-293T cells cultured in Opti-MEM was higher compared to cells cultured in EV-DEP DMEM but this was not significant, likely due to inter-experiment variability (Supplementary Figure 3F). Western blotting shows that HEK-293T cells release more CD9 and CD63 positive EVs/1 million cells when cultured in Opti-MEM as compared to EV-DEP DMEM (Supplementary Figure 3G, H& I). Overall, the characterisation of EVs isolated from MDA-MB-231 and HEK-293T cells supports our findings from the analysis of EVs released into conditioned media by MDA-MB-231-zsGreen and HEK-293T-myr-palm-mNeonGreen(2x) cells (Figure 1): cells cultured in Opti-MEM release significantly more EVs than in EV-DEP DMEM.

### The proteomic composition of Opti-MEM and EV-DEP DMEM derived EVs is largely unchanged, but EV-DEP DMEM EVs contain significantly more contaminating histone and bovine proteins

Cells cultured in Opti-MEM release more EVs; we next wanted to ascertain whether the proteomic profile of EVs is also changed. To do this, 4 µg of EVs from MDA-MB-231 cells cultured in either Opti-MEM or EV-DEP DMEM were subjected to proteomic analysis. GO-analysis of the mass spectrometry show that the proteins arise from cellular compartments synonymous with EVs biogenesis such as membrane, vesicle, cytosol, extracellular space, endosome, and membrane-enclosed lumen (WebGestalt, 2024). (Supplementary Figure 5). Furthermore, the majority of the top 100 identified proteins in over three thousand EV mass spectrometry experiments (Vesiclepedia, 2025) were also found in our EV mass spectrometry protein identifications (Supplementary table 1). A comparison of the protein composition of EVs derived from Opti-MEM vs EV-DEP DMEM culture conditions shows that the majority of proteins identified were similar (2212 out of 2285), with only a small number of unique proteins found in EVs from cells cultured in Opti-MEM (n=12) or DMEM (n=61) (Figure 3A) (Venny 2025). Further analysis by Mass Dynamics shows that that there were only two proteins significantly elevated in EVs from cells cultured in Opti-MEM as compared to 35 proteins significantly elevated in EVs from cells cultured in EV-DEP DMEM (Figure 3B, Supplementary table 2). Further assessment of the proteins that were significantly elevated in EVs derived from EV-DEP DMEM culture conditions revealed a trend: the elevated proteins belonged to one of two groups – nucleus-derived or protein corona derived (Table 1). For example, there were 12 histone proteins significantly elevated in EV-DEP DMEM EVs as well as four other nuclear proteins. Additionally, there were 14 serum-derived bovine proteins (including albumin) significantly elevated in DMEM EVs, as well as five other ECM and protease related proteins, all of which likely contribute to the EV protein corona.

**Figure 3:**
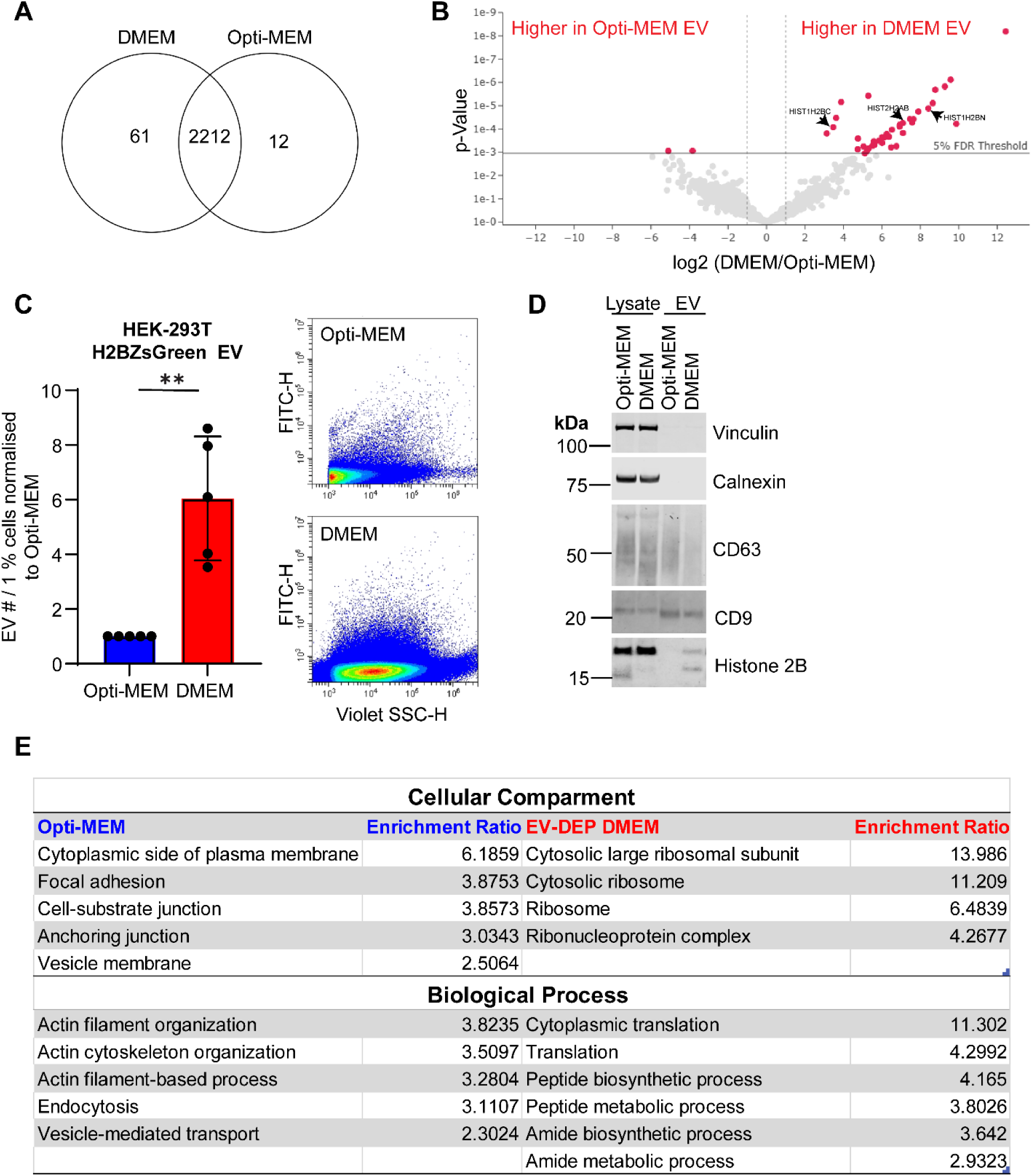
EVs released by MDA-MB-231 cells in EV-DEP DMEM contain more histones and bovine proteins than those in Opti-MEM. Proteins in EVs isolated from MDA-MB-231 cells cultured in EV-DEP DMEM or Opti-MEM were analysed by mass spectrometry. N=3. Common and unique protein IDs are shown in Venn diagram – Venny 2025 (A). Mass Dynamics analysis of LFQ intensities showed only two proteins were significantly elevated in Opti-MEM EVs, compared to 38 proteins, including histones, in EVs from cells cultured in EV-DEP DMEM (B). HEK-293T cells were transiently transfected with Histone2B-ZsGreen and the number of Histone2B positive EVs released by cells in Opti-MEM vs EV-DEP DMEM analysed by nanoflow cytometry with representative flow plots. N=5. ± SEM. One sample T-test (C). To validate the mass spectrometry results, the presence of EV markers CD9 and CD63, contaminant calnexin, and Histone2B in 2 µg EVs from MDA-MB-231 cells cultured in Opti-MEM or EV-DEP DMEM was analysed by western blot. N=3. (D). WebGestalt was used to analyse the top 70 % of proteins enriched in Opti-MEM compared to EV-DEP DMEM, and EV-DEP DMEM compared to Opti-MEM. The enrichment of proteins associated with cellular compartments and biological processes are represented (E).

**Table 1:** Mass spectrometry analysis of proteins that are significantly elevated in MDA-MB-231 EVs from cells cultured in EV-DEP DMEM versus Opti-MEM.

| Significantly higher in: | Protein name: | Protein category: |
| --- | --- | --- |
| EV-DEP DMEM | Transmembrane protein 109 | Nucleus/DNA damage response |
|  | Histone H2B type 1-N | Nucleus |
|  | Histone H2A type 2-B | Nucleus |
|  | UHRF1-binding protein 1 | Nucleus |
|  | Histone H2B type 2-E | Nucleus |
|  | Histone H3.1 | Nucleus |
|  | Histone H2B type 1-C/E/F/G/I | Nucleus |
|  | Histone H4 | Nucleus |
|  | Histone H2A type 1-D | Nucleus |
|  | Histone H1.2 | Nucleus |
|  | Histone H2AX | Nucleus |
|  | Histone H2A.V;Histone H2A.Z | Nucleus |
|  | U1 small nuclear ribonucleoprotein 70 kDa | Nucleus |
|  | Histone H1.0;Histone H1.0, N-terminally processed | Nucleus |
|  | Thymidylate synthase | Nucleus |
|  | Core histone macro-H2A.1 | Nucleus |
|  | Thioredoxin-related transmembrane protein 1 | Vesicular trafficking/golgi/ER |
|  | Vesicle transport protein GOT1B | Vesicular trafficking/golgi/ER |
|  | Transmembrane emp24 domain-containing protein 10 | Vesicular trafficking/golgi/ER |
|  | Serine protease HTRA1 | Protease/protease inhibitor |
|  | Kunitz-type protease inhibitor 2 | Protease/protease inhibitor |
|  | Serglycin | ECM |
|  | Thrombospondin-1 | ECM |
|  | Protein CYR61 | ECM |
|  | Hemoglobin fetal subunit beta | BOVINE |
|  | Serpin A3 | BOVINE |
|  | Serpin A3-7-like precursor | BOVINE |
|  | Fetuin-B | BOVINE |
|  | Inter-alpha-trypsin inhibitor HC2 component homolog | BOVINE |
|  | Vitronectin | BOVINE |
|  | Apolipoprotein E | BOVINE |
|  | Alpha-1-antiproteinase | BOVINE |
|  | Clusterin | BOVINE |
|  | Immunoglobulin heavy constant mu | BOVINE |
|  | Alpha-2-HS-glycoprotein | BOVINE |
|  | Albumin | BOVINE |
|  | Hemoglobin subunit alpha | BOVINE |
|  | Prothrombin | BOVINE |
| Opti-MEM | Ubiquitin-fold modifier-conjugating enzyme 1 | Ubiquitin pathway |
|  | Carbonic anhydrase 12 | Enzyme |

To validate the mass spectrometry findings, we evaluated Histone2B levels in EV preparations from HEK-293T and MDA-MB-231 cell lines. Histone2B-ZsGreen was overexpressed in HEK-293T cells and 24 hours later 20,000 cells were plated in each well of a 96 well plate in either Opti-MEM or EV-DEP DMEM media. After 24 hours of cells conditioning the media, EV and small particle release was quantified by nanoflow cytometry. Confirming the mass spectrometry results, more Histone2B-ZsGreen positive events were recorded when HEK-293T cells were cultured in EV-DEP DMEM as compared to Opti-MEM (Figure 3C). It is worth noting that the Histone2B-ZsGreen positive particles were not sensitive to detergent lysis suggesting that these particles are not EVs nor histones associated with EVs (Supplementary Figure 6). Next, we isolated EVs from MDA-MB-231 cells cultured in Opti-MEM or EV-DEP DMEM, loaded by protein concentration, and assessed for EV abundance by western blot. Increased levels of CD9 and CD63 was identified for the Opti-MEM culture condition compared to EV-DEP DMEM, whereas cells cultured in EV-DEP DMEM released more Histone2B positive particles (Figure 3D). This further supports the increased EV production by cells cultured in OptiMEM and validates the mass spectrometry data (Figure 3D). Overall, the mass spectrometry analysis shows that the proteome composition of EVs and small particles from cells cultured in Opti-MEM versus EV-DEP DMEM is largely unchanged except for the presence of histones and bovine proteins in DMEM-derived EVs, these are likely soluble proteins making up part of the EV protein corona or small particle aggregates that co-isolated with EVs during ultracentrifugation.

While the overall proteomic profile of EVs isolated from cells cultured in Opti-MEM or EV-DEP DMEM is similar, (aside the bovine proteins and histones), there were notable changes in protein abundance between the two conditions. The top 70 % of proteins enriched in Opti-MEM as compared to DMEM (and vice-versa) were subjected to gene enrichment analysis and the cellular compartments and biological processes in which the identified proteins cluster assessed. While proteins identified in Opti-MEM were found to reside in the cytoplasmic side of the plasma membrane, focal adhesions, cell-substrate junctions, anchoring junctions, and vesicle membranes, proteins enriched in EV-DEP DMEM EVs were found to more commonly associate with ribosomal structures (Figure 3E). Furthermore, proteins enriched in Opti-MEM EVs were found to be involved in actin cytoskeleton dynamics, endocytosis, and vesicle mediated transport, compared to proteins enriched in EV-DEP DMEM EVs which were found to be involved in processes such as protein translation and peptide/amide metabolic processes. Overall, the proteins most enriched in Opti-MEM EVs, as compared to EV-DEP DMEM EVs, were associated with cellular localisation and biological processes that are more traditionally associated with EV biogenesis and release pathways.

### The mechanism of EV release by MDA-MB-231 cells differs depending on whether cells are cultured in Opti-MEM or EV-DEP DMEM

To evaluate EV release mechanisms, MDA-MB-231 cells were engineered to stably express a PalmGRET reporter construct. The PalmGRET construct contains the growth-associated protein 43 (GAP43) palmitoylation sequence anchored to a GFP-NLuc BRET reporter which has been previously demonstrated to enrich the GFP-NLuc BRET reporter in small-extracellular vesicles^19^ (Figure 4A). The enrichment and increased brightness of EVs from MDA-MB-231-PalmGRET cells provides a more robust method to quantify both large (10,000 x g pellet) and small (remaining EVs in conditioned media) EV populations by nanoflow cytometry. Figure 4B & E shows that similar to the zsGreen and myr-palm-mNeonGreen(2x) systems, PalmGRET enriched small EVs were released in larger numbers when MDA-MB-231 cells were cultured in OptiMEM as compared to DMEM. Quantification of large EV (10,000 x g pellet) production from MDA-MB-231-PalmGRET cells cultured in Opti-MEM or EV-DEP DMEM identified a significant reduction in the number of large EVs (10,000 x g pellet) in the Opti-MEM culture condition (Figure 4B). Importantly, large and small PalmGRET positive EVs were lysed by 1:1 dilution in 1 % Triton-X indicating that bonafide GFP containing EVs are being analysed (Figure 4C & D).

**Figure 4:**
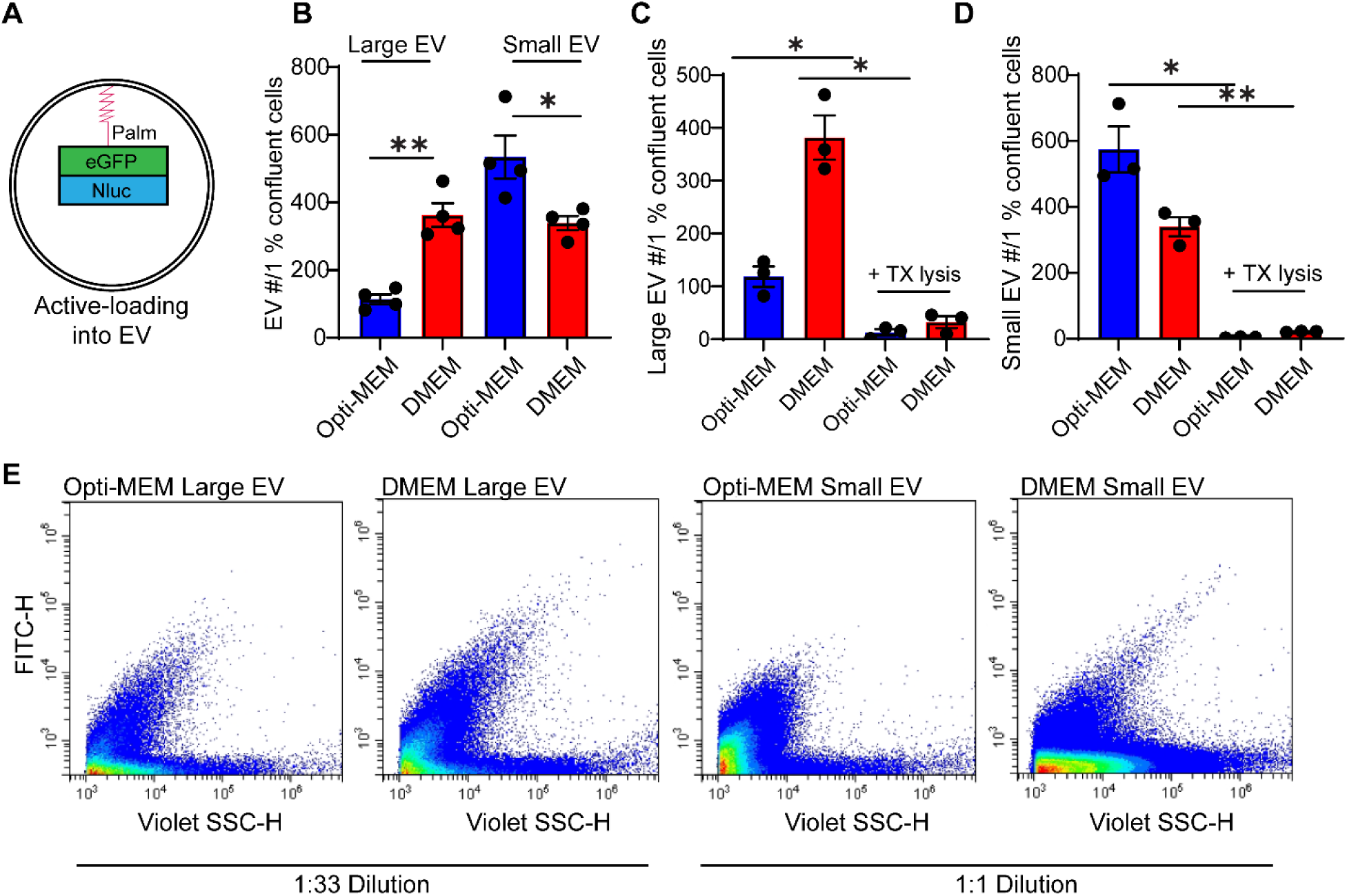
MDA-MB-231 cells stably expressing PalmGRET plasmid release more GFP positive small EVs when cultured in Opti-MEM as compared to DMEM. MDA-MB-231 cells stably expressing PalmGRET (A) were plated in 12 well plates, 24 hours later media was refreshed with EV-DEP DMEM or Opti-MEM and cells were allowed to condition media for 24 hours. Large (10,000 x g pellet) and small (remaining EVs in conditioned media) EVs were measured using nanoflow cytometry (B). To test lysis efficiency, small or large EVs were diluted 1:1 in 1 % Triton-X for 30 minutes before being measured by nanoflow cytometry (C & D). Representative flow plots showing large and small EV populations from MDA-MB-231 PalmGRET cells growing in Opti-MEM or EV-DEP DMEM (E).

As the Palm-GRET system supported quantification of both small and large EVs in small volumes of conditioned media, we evaluated possible changes to the mechanism of EV release dependent on the cell culture medium. To begin an assessment of EV-release mechanisms, established EV modulators were tested in different mediums. Bafilomycin (160 nm) was used to stimulate EV release, while sphingomyelinase inhibitor GW4869 (5 µM) and ROCK kinase inhibitor Y-27632 (5 µm) were used to perturb EV release. Analysis of large and small EVs released from treated cells revealed differences in EV release mechanisms dependent upon the medium. Bafilomycin treatment was found to effectively increase large and small EV release from cells cultured in Opti-MEM and EV-DEP DMEM compared to DMSO control (Figure 5A & B). It is also noteworthy that DMSO (1:1000) decreased small EV release in EV-DEP DMEM but not in Opti-MEM, further emphasising the use of appropriate control compounds in pharmacological studies. Furthermore, inhibition of the sphingomyelinase pathway (GW4869) in cells cultured in Opti-MEM, decreased large EV release by 52 % and small EV release by 59 % compared to DMSO control (Figure 5A & C), compared to 46 % and 31 % in large and small EVs released from cells cultured in EV-DEP DMEM (Figure 5B & D). Cells in EV-DEP DMEM were more dependent on ROCK kinases for the release of membrane derived ectosomes as treatment with Y-27632 reduced the number of large EVs by 46 % and small EVs by 45 %) (Figure 5B & F) whereas in Opti-MEM, Y-27632 reduced the number of large EVs by 24 % and small EVs by 15 % (Figure 5A & E).

**Figure 5:**
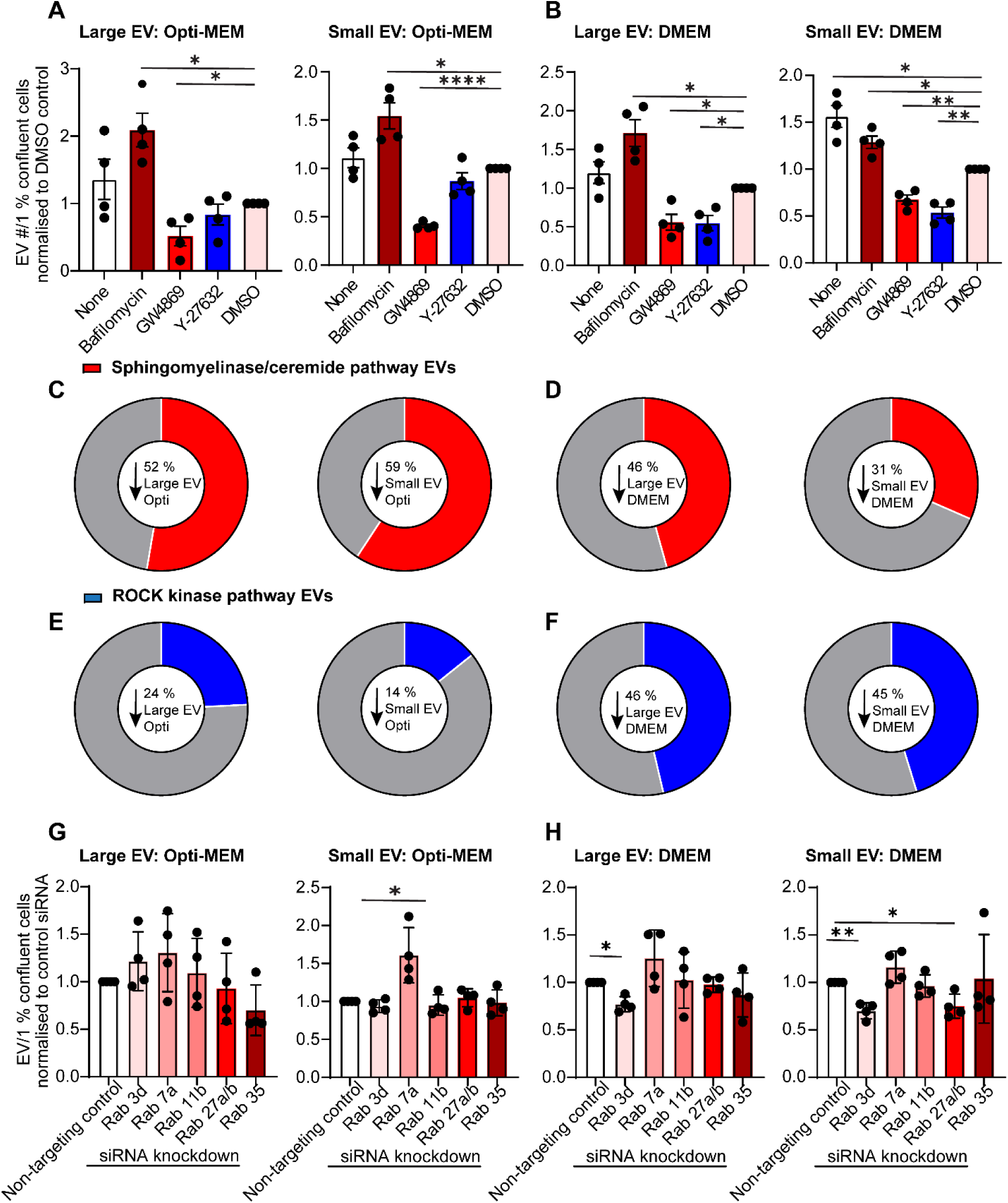
EVs released by MDA-MB-231-PalmGRET cells in Opti-MEM use predominantly the sphingomyelinase pathway, whereas cells in EV-DEP DMEM rely more on ROCK kinases and RabGTPases for EV release. MDA-MB-231 PalmGRET cells were plated into each well of a 12 well plate, 24 hours later cells were washed 2x with PBS and replenished with 1.5 ml either Opti-MEM (A) or EV-DEP DMEM (B) containing Bafilomycin (160 nm), GW4869 (5 µm), Y27632 (5 µm). After 24 hours, conditioned media was spun 300 x g 10 min, 2500 x g 15 min, and 10,000 x g 30 mins to isolate large EVs, the remaining EVs in the supernatant were small EVs. Large and small EVs were quantified by nanoscale flow cytometry. The percentage decrease in release of large and small EVs by cells cultured in Opti-MEM or EV-DEP DMEM after treatment with GW4869 (C & D) or Y27632 (E & F) compared to DMSO control was represented in pie charts. Finally, siRNA to Rab 3d, 7a, 11a, 27a&b, and 35 was transfected into MDA-MB-231-PalmGRET cells. Twenty four hours after transfection cells were replated and 24 hours later media was refreshed to 1.5 ml of either Opti-MEM or EV-DEP DMEM. Cells conditioned media for 24 hours and upon harvest, spun 300 x g 10 min, 2500 x g 15 min, and 10,000 x g 30 mins. Small EVs (remaining in conditioned media) and large EVs (from the 10,000 x g pellet) were analysed by nanoflow cytometry from cells incubated in Opti-MEM (G) or EV-DEP DMEM (H).

To determine which RabGTPases are necessary for small EV release in Opti-MEM and EV-DEP DMEM, MDA-MB-231-PalmGRET cells were plated in a 12 well plate and transfected with siRNAs to Rab GTPases either known to be involved in EV biogenesis or identified in the proteomics data (Rab 3d, 7a, 11b, 27a&b, & 35). Media was conditioned for 48-72 hours post-transfection and analysed for changes in the number of EVs released. We found that MDA-MB-231-PalmGRET cells cultured in EV-DEP DMEM relied on Rab3d for small and large EV release, and Rab27a and Rab27b for small EV release (Figure 5H). Conversely, cells cultured in Opti-MEM did not rely on any of the Rab GTPases tested (Rab3d, 7a, 11b, 27a&b, 35) for EV release (Figure 5G).

## Discussion

The selection of cell culture media is an important experimental variable that can impact EV yield, cargo, purity, and physiology. Due to the absence of bovine-derived contaminants, lack of batch to batch variability, decreased immunogenicity, and increased compliance with regulatory standards, serum-free and animal-component free medias are desirable options for the cell culture and production of cell-based and cell-free biologic therapeutics such as EVs^25,26^. Here, we aimed to determine what impact serum free medias had upon EV biogenesis, release, and purity.

We demonstrate that cells cultured in Opti-MEM released a larger number of EVs than in EV-DEP DMEM, which is beneficial for experiments that require large EV yields. Conversely, cells cultured in EV-DEP DMEM released fewer EVs and these EV preparations are rich in histones and bovine proteins, both of which have the potential to be immunomodulatory^12,13,27–29^. Our study aligns with others that have investigated the use of serum-free media for EV production. For example, a neuroblastoma cell line grown in serum-free conditions released significantly more EVs with an altered EV proteome^30^. A second publication reported that HEK-293 cells in serum-free Opti-MEM released more EVs in a ceremide-dependent manner and these EVs had altered transcriptional profiles^31^. Our data is in-line with these studies, and we report an increase in EV release from MDA-MB-231 and HEK-293T cells in Opti-MEM compared to EV-DEP DMEM. In addition, EV release from MDA-MB-231 cells in Opti-MEM were highly dependent upon the ceremide biogenesis pathway. The sphingomyelinase inhibitor GW4869 decreased small EV release from cells in Opti-MEM by 59 % compared to just 31 % from cells in EV-DEP DMEM. Further, we find that cells cultured in EV-DEP DMEM have more dependence upon ROCK kinases for membrane-derived EV biogenesis. The ROCK kinase inhibitor decreased small EV release in EV-DEP DMEM by 45% compared to only 15% in Opti-MEM. This suggests that EV-DEP DMEM supports the release of more membrane derived ectosomes as compared to cells in Opti-MEM.

Cells in EV-DEP DMEM did demonstrate some sensitivity to the sphingomyelinase inhibitor GW4869 suggesting that they still do release MVB-derived exosomes, but given our results with the ROCK kinase inhibitor, ectosomes may be a more prominent EV population under this culture condition in comparison to Opti-MEM. Instead of being reliant upon the sphingomyelinase pathway like cells cultured in Opti-MEM, cells in EV-DEP DMEM use multiple mechanisms for MVB-derived exosome release including the sphingomyelinase pathway, and Rab3d and Rab27a/b. Rab27a/b is a well characterised and established mechanism of exosome release thought to be essential for MVB transport and docking to the membrane^32^. It is interesting that a traditional and well characterised route of exosome biogenesis is differentially utilised under different cell culture conditions and may not always be the dominant route depending upon cell culture model, conditions, media, and supplementation.

The dependence of MDA-MB-231 MVB-derived exosome release on Rab3d is supported by a previous publications which describe the importance of Rab3d in MVB transport and docking^33^ and pro-metastatic EV release in MDA-MB-231 cancer cells^34^. As Rab3d is important in EV-DEP DMEM EV release but has no role in cells cultured in Opti-MEM, it would be interesting to determine whether the EVs released from cells in the two different medias, exert differing pro-metastatic effects upon recipient cells. It may be important to consider using a media composition that reflects the nutrient availability of individual cancer types in-vivo to ensure that EVs are released using the molecular pathways that would be active in-vivo, and therefore their phenotypic properties are reflective of what would be found in physiology. Opti-MEM effectively creates an environment of low nutrient availability which is the norm for some solid tumors^35,36^. However, some cancer cells such as those on the tumor edge or at metastatic sites^37^ may have different nutrient exposures or even higher nutrient availability and therefore, EV-DEP DMEM may be a better choice of media for culture. The physiological nutrient availability of the studied cell type should be taken into consideration when selecting a media for use in EV biogenesis, mechanism. and function studies.

The proteome composition of EVs isolated from Opti-MEM and EV-DEP DMEM cells were remarkably similar, except for two protein groups: Histone and bovine proteins were elevated in EVs from EV-DEP DMEM cells. The presence of bovine byproducts (EVs, proteins, lipoproteins, nucleic acids, and lipids) in EV preparations can cause several problems in downstream applications. For example, bovine component contamination of EVs has been found to contribute to changes in cell proliferation^10,38–40^, migration^11^, immune cell suppression^12,13^, differentiation^38,39^, and EV release^30,31^. Additionally, it is known that serum-derived components unintentionally delivered as part of a cell therapy protocol can initiate an immune response in some individuals, with anti-bovine antibodies being detected in the blood of some patients^28,29^. This portrays the importance of using media devoid of serum-derived bovine proteins for therapeutic development and clinical applications.

Histones have been widely reported to be present in EV preparations for many years, with some uncertainty surrounding whether they are co-isolated contaminants or are selectively sorted and integrated into EVs/EV membranes as carriers of DNA^41–44^. A recent study found that cell stress conditions such as heat challenge, inflammation, or hypoxia increased the secretion of histone-containing EVs produced through the multivesicular body pathway^43^. Our mass spectrometry data supports the presence of histones in EV preparations, but histone-positive particles were not sensitive to detergent treatment suggesting that they were not EV-associated or encapsulated. Instead, the histones are likely a soluble protein aggregation either co-isolated with EVs or associated with EV membranes as a protein corona component. Our previous publication showed that soluble proteins such as mammaglobin-A can associate with EVs, but upon EV lysis, only CD9-mammaglobin-A positive EV events lysed and the larger population of mammaglobin-A, CD9 negative, events did not lyse^45^. A proportion of the histone positive events may be EV associated which could be tested by including an EV marker such as CD9. Although the pathophysiological consequence of histones in EV preparations has not yet been fully characterised, it is known that the presence of extracellular/circulating histones in cases of injury and trauma can cause recipient cell stress, immune cell stimulation, and inflammatory responses^27^. Therefore, cell media choice and the impact it may have upon histone presence in EV preparations should be carefully considered and characterised if the EVs are to be used in vivo, in therapeutic development, or in immune cell assays.

Overall, cells in Opti-MEM produce a higher yield of purer (histone and bovine protein depleted) sphingomyelinase MVB-derived EVs, whereas EV-DEP DMEM produces a lower yield of ROCK dependent membrane-derived ectosomes and sphingomyelinase, Rab3d, Rab27a&b, dependent MVB-derived EVs that are rich in immunostimulatory histones and bovine proteins. The data indicates that EVs derived from cells grown in EV-DEP DMEM may be more physiologically relevant when studying mechanisms and functions of nutrient rich cancer cells. However, EVs-derived from cells grown in Opti-MEM would be more desirable for use in the study of: 1) traditionally nutrient deprived cancer cells from solid tumors for example, and 2) for immune assays or therapeutic applications due to the higher yield of EVs that contain fewer immunostimulatory histones and bovine proteins. It would still be imperative to identify if other nucleic acids, metabolites, or lipid components are altered in EVs from cells cultured in Opti-MEM before use in therapeutic applications to ensure that no undesirable cargoes could be transferred to recipient cells by the EVs that could cause cell stress, toxicity, or carcinogenicity.

### Conclusion

So DMEM, or Opti-MEM, for EV isolation and experimentation? There is no one correct answer that suits every experiment. The choice of media should be carefully considered when designing each individual experiment, and the media choice should be based on considerations for experimental outcomes, yield, desired EV sub-type, and the intended downstream applications.

## Acknowledgements

Mass Spectrometry analysis was performed in the Proteomics and Metabolomics Core Facility – at the Michael Smith Laboratories and the Life Sciences at the University of British Columbia supported by Jason Rogalski and Renata Moravcova. Electron microscopy was done at the University of British Columbia Bioimaging Facility.

## Declaration of Interest Statement

The authors report no conflict of interest. The project was funded by a Natural Sciences and Engineering Research Council of Canada (NSERC) Discovery Grant (RGPIN-2020-04641) awarded to KCW.

## Data Sharing

All data, materials and methods are available upon request. The raw mass spectrometry data is available on ProteomeXchange via the MassIVE (Mass Spectrometry Interactive Virtual Environment) partner repository. Data set identifier: PXD068319.

## Notes

### Competing Interest Statement

The authors have declared no competing interest.

### Summary of Updates

The abstract was not amended on the last resubmission to reflect the new data. The abstract now reads to inform on the new data in figure 4 and 5.

