## Supplemental Figures for "DMEM, or Opti-MEM, that is the Question: An Important Consideration for Extracellular Vesicle Isolation and their Downstream Applications"

**Supporting Data**

Supplementary Figures 1 – 7

Supplementary Tables 1 & 2

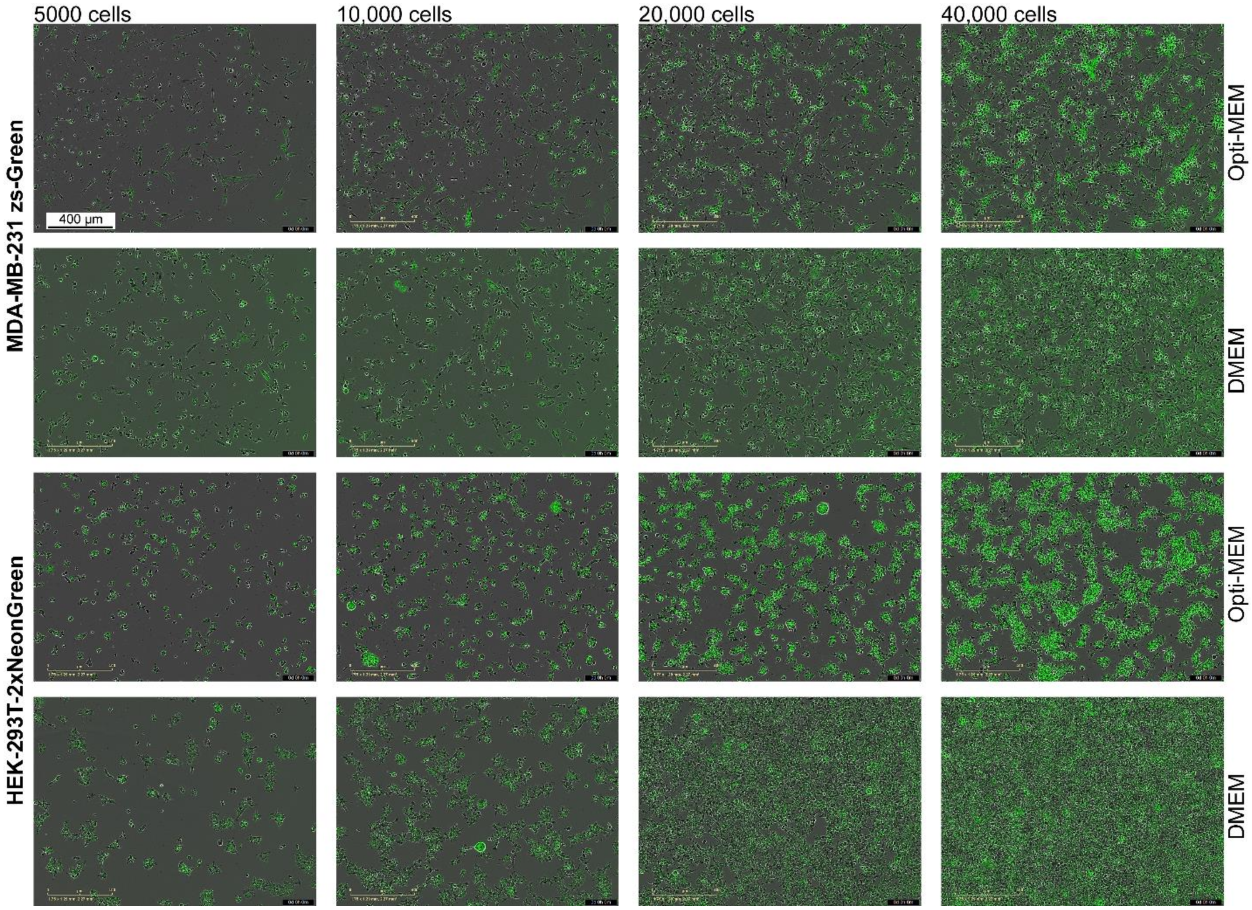


**Supplementary Figure 1: Cell morphology of 231-zsGreen cells and HEK-293T-mNeonGreen(2x) cells cultured at different confluences in Opti-MEM or 10 % FBS DMEM cell culture media.** 5,000, 10,000, 20,000 or 40,000 MDA-MB-231-zsGreen cells or HEK-293T-Myr-Palm-mNeonGreen(2x) cells were plated per well in a 96 well plate in Opti-MEM or 10 % serum supplemented DMEM for 48 hours to condition media. At 48 hours the IncuCyte-Sx5 was used to image and analyse cell confluence, zsGreen/NeonGreen(2x) expression and cell morphology. Representative images chosen from one experiment of three.

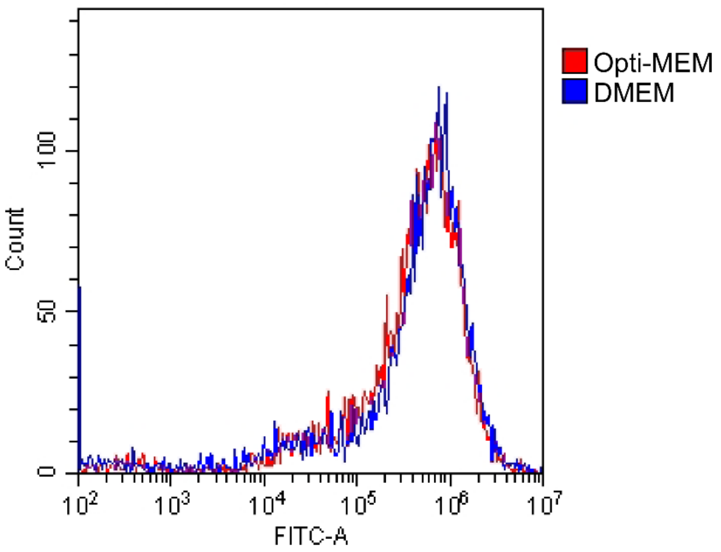


**Supplementary Figure 2: MDA-MB-231-zsGreen cells have similar zsGreen fluorescent intensities when cultured in both Opti-MEM and DMEM.** MDA-MB-231-zsGreen cells were grown in either Opti-MEM or 10 % serum supplemented DMEM for 24 hours before trypsinisation and analysis of their brightness using flow cytometry (CytoFLEX-S, Beckman).

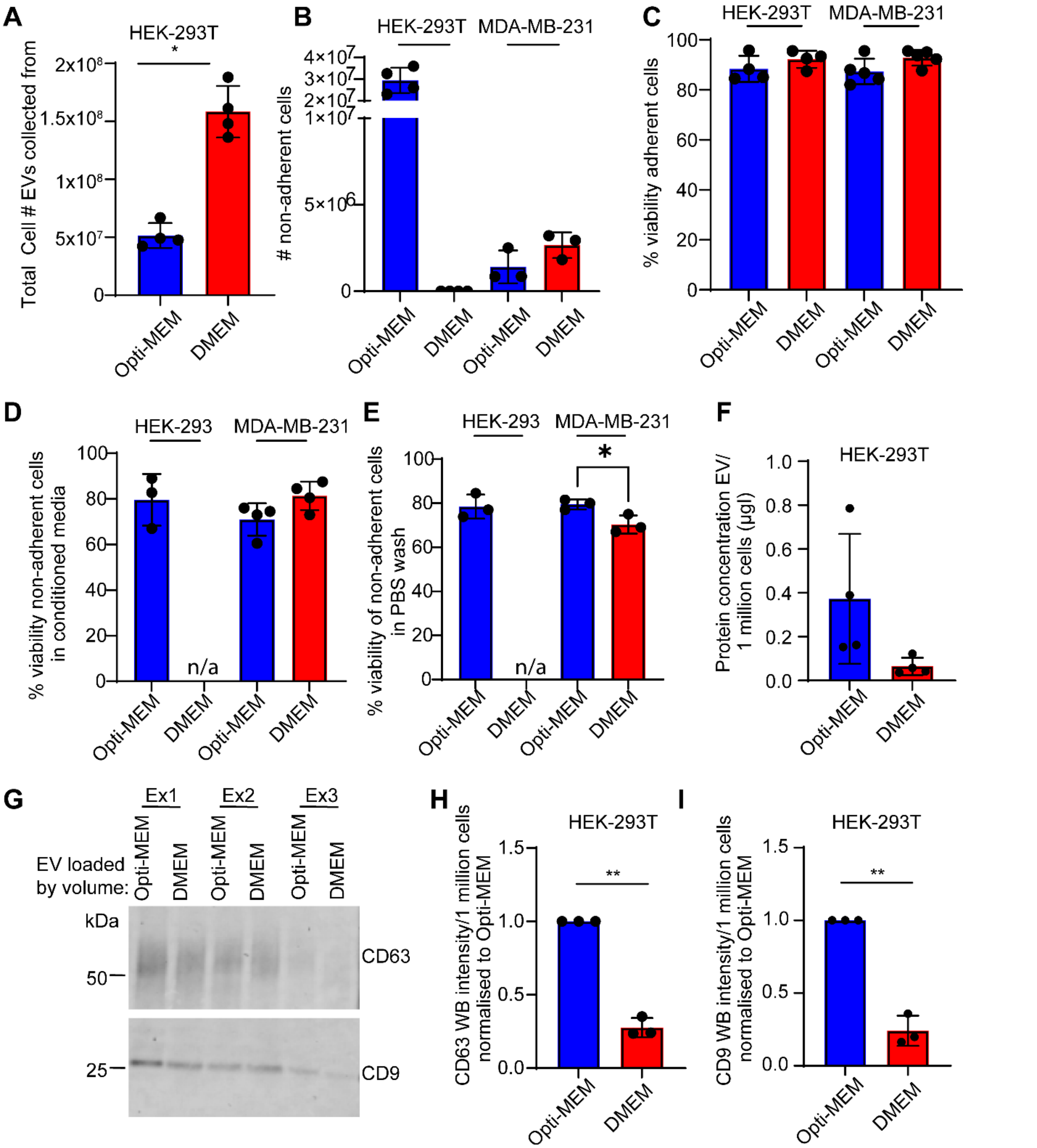


**Supplementary figure 3: HEK-293T-zsGreen cells release more EVs when cultured in Opti-MEM as compared to EV-DEP DMEM media.** HEK-293T-zsGreen cells were cultured in Opti-MEM or EV-DEP DMEM for 48 hours before EVs were harvested using differential ultracentrifugation. (A) The total cell number and (B) the number of non-adherent floating cells from which EVs were harvested from was recorded alongside the cell viability of adherent and non-adherent cells (C, D, & E). (F) The protein concentration of EVs released by 1 million cells was ascertained using the micro-BCA assay. (G) HEK-293T EVs from three experiments were western blotted for EV markers CD9 and CD63, the relative abundance of each marker in EVs from cells cultured in Opti-MEM versus DMEM was calculated using densitometry, normalized to cell number and displayed in graphs (H) CD63, and (I) CD9.

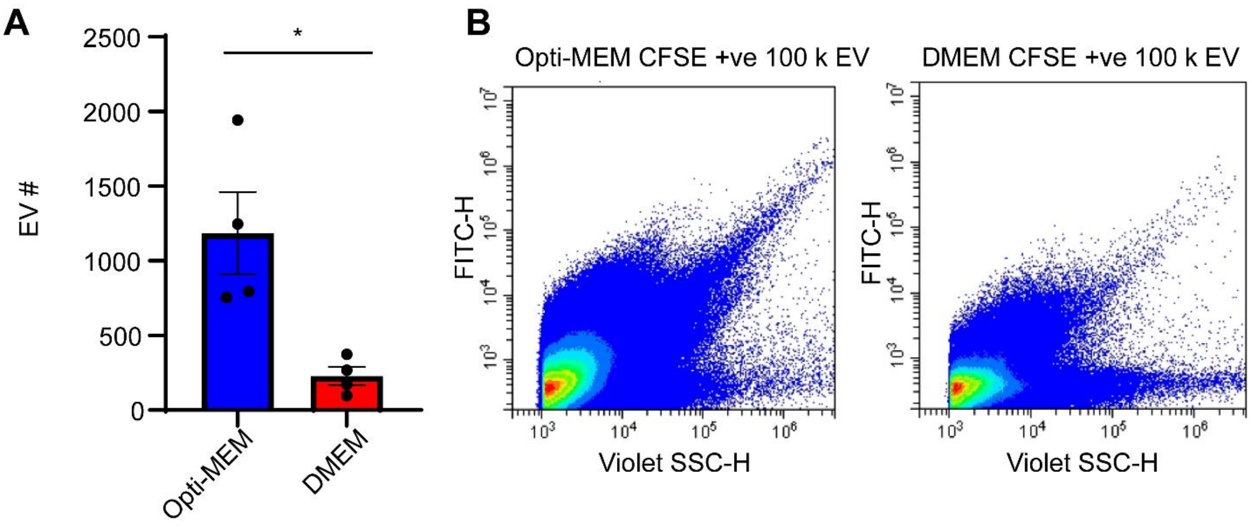


**Supplementary Figure 4: MDA-MB-231 cells cultured in Opti-MEM release more CFSE positive 100,000 x g pellet EVs than cells cultured in DMEM.** (A) MDA-MB-231 cells were cultured in Opti-MEM or EV-DEP DMEM for 48 hours before EVs were harvested from conditioned media by differential ultracentrifugation. The 100,000 x g pellet containing small EVs were stained green using CFSE (1:2000) for 10 minutes at 37 °C before analysing EV events using nanoflow cytometry (1:1000 dilution). N=4. ± SEM. (B) Representative flow plot images.

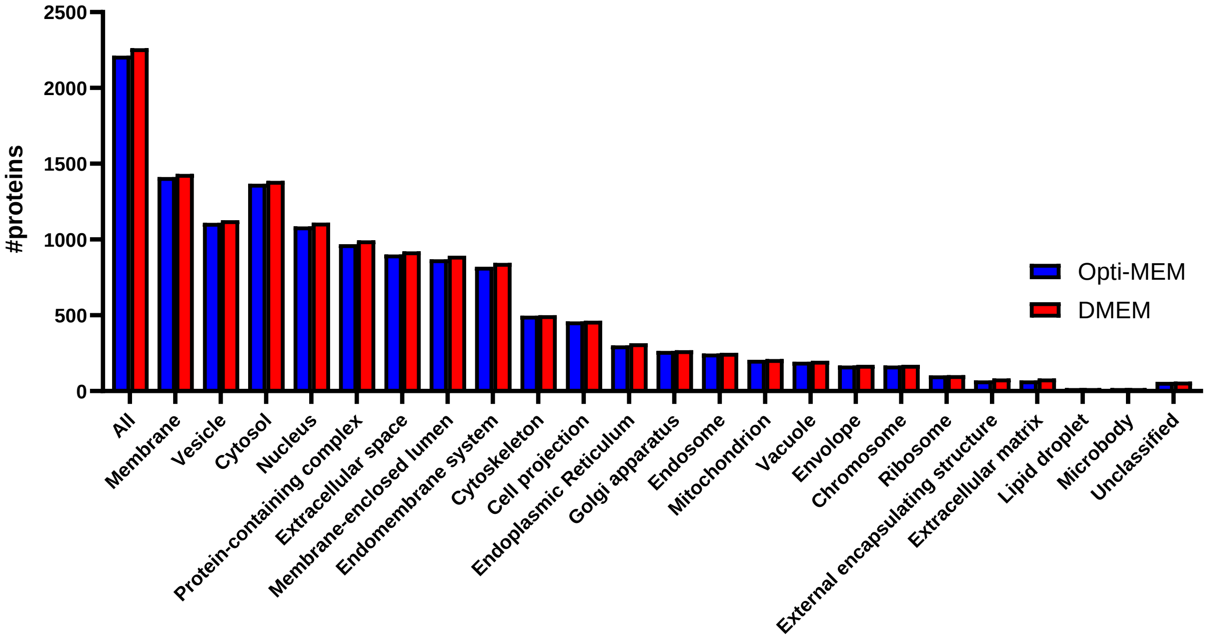


**Supplementary Figure 5: GO-analysis of proteins identified in mass spectrometry of MDA-MB-231 EVs.** WebGestalt (2024) was used for GO-analysis of the EV proteins identified to be present in preparations from MDA-MB-231 cells cultured in either Opti-MEM or EV-DEP DMEM. The cellular compartments in which identified EV proteins reside are shown.

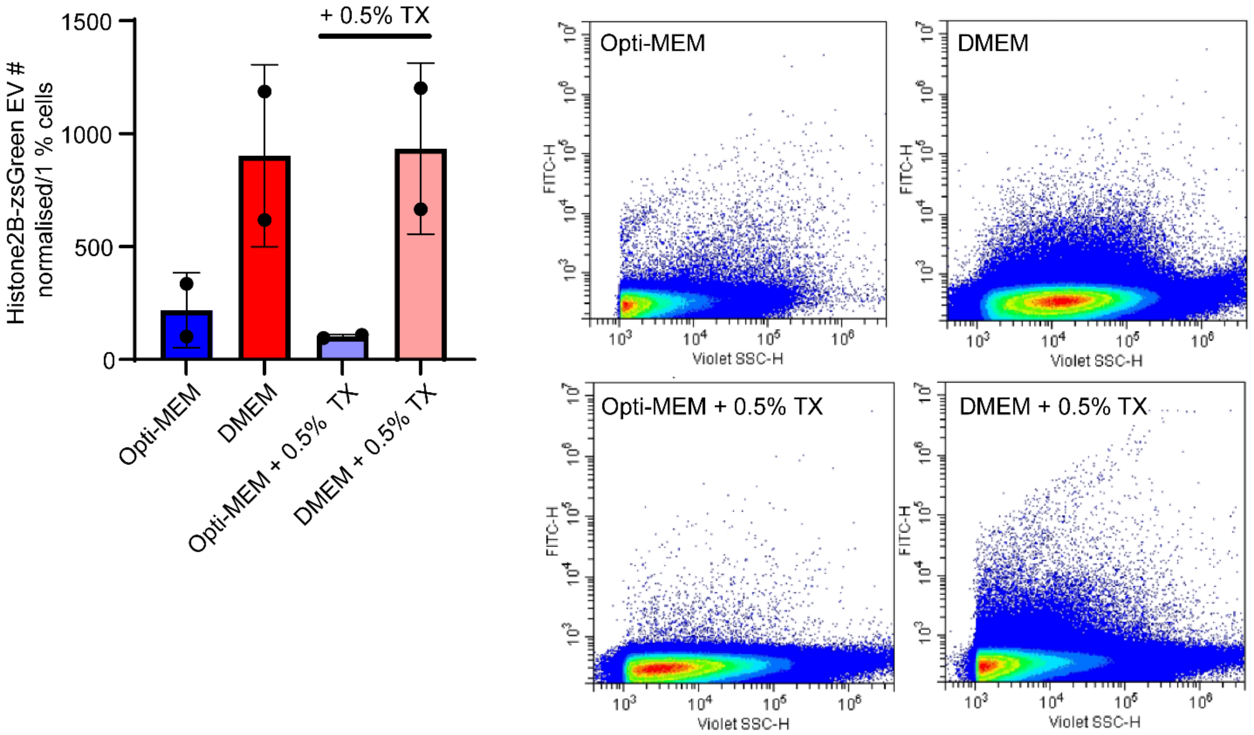


**Supplementary Figure 6: Histone2B-zsGreen positive HEK-293T cell derived EVs do not efficiently lyse.** HEK-293T cells were transiently transfected with H2B-zsGreen plasmid and 24 hours later plated in a 96 well plate and allowed to condition media for a further 24 hours. The number of H2B-zsGreen positive EVs was analysed in conditioned media diluted 1:1 with PBS by nanoflow cytometry before and after lysis for 10 minutes by adding 1% Triton-X 1:1 with conditioned media:PBS solution to a final 0.5 % concentration. EV number after lysis multiplied 2x to correct for dilution. Representative flow plots shown. N=2.

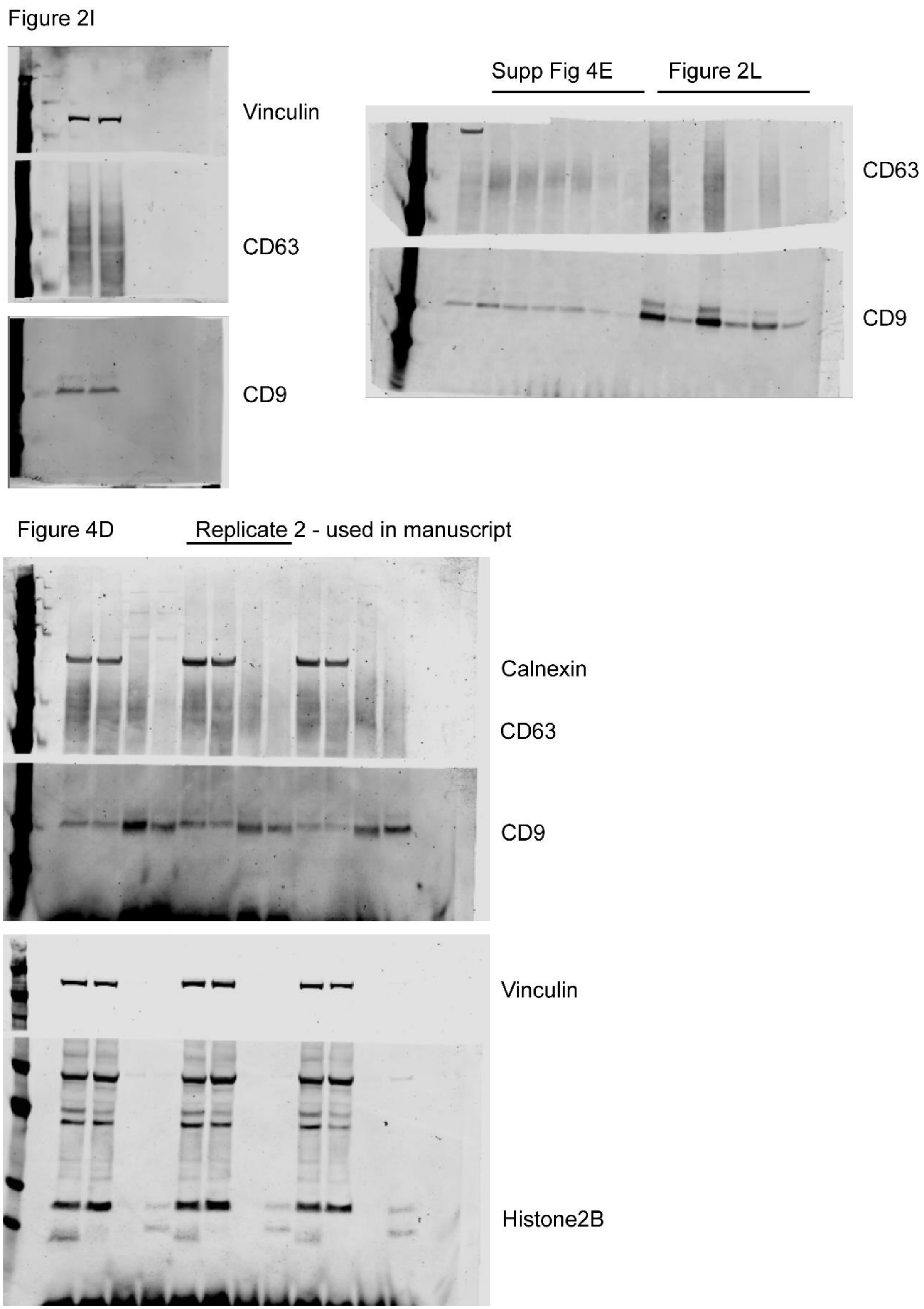


**Supplementary Figure 7: Full length Western blots.**

**Supplementary Table 1: The 100 most commonly identified EV proteins in mass spectrometry experiments (Vesiclepedia 2025) also identified in in MDA-MB-231 cell derived EVs after culture in Opti-MEM or EV-DEP DMEM. Blue bracket = proteins elevated in Opti-MEM EVs. Red bracket = proteins elevated in DMEM EVs.**

| Protein | Opti-MEM | DMEM | Opti-MEM/DMEM ratio |
|---|---|---|---|
| A2M | Y | N | n/a |
| CD63 | N | N | n/a |
| CD81 | N | N | n/a |
| HIST1H4A | N | N | n/a |
| C3 | Y | Y | 24.26 |
| FN1 | Y | Y | 7.88 |
| LGALS3BP | Y | Y | 2.68 |
| CLTC | Y | Y | 2.66 |
| ALB | Y | Y | 2.35 |
| AHCY | Y | Y | 1.68 |
| MFGE8 | Y | Y | 1.67 |
| EEF1G | Y | Y | 1.64 |
| EEF1A1 | Y | Y | 1.60 |
| CCT3 | Y | Y | 1.59 |
| IQGAP1 | Y | Y | 1.54 |
| CCT4 | Y | Y | 1.51 |
| HSPA5 | Y | Y | 1.49 |
| FLNA | Y | Y | 1.48 |
| CCT6A | Y | Y | 1.48 |
| TCP1 | Y | Y | 1.45 |
| FLOT2 | Y | Y | 1.42 |
| RAN | Y | Y | 1.41 |
| SDCBP | Y | Y | 1.37 |
| CCT5 | Y | Y | 1.34 |
| EEF2 | Y | Y | 1.30 |
| ANXA2 | Y | Y | 1.28 |
| CCT2 | Y | Y | 1.27 |
| CCT8 | Y | Y | 1.26 |
| HSP90AB1 | Y | Y | 1.22 |
| EIF4A1 | Y | Y | 1.21 |
| LDHA | Y | Y | 1.20 |
| KPNB1 | Y | Y | 1.16 |
| HSP90AA1 | Y | Y | 1.13 |
| LDHB | Y | Y | 1.12 |
| MYH9 | Y | Y | 1.11 |
| VCP | Y | Y | 1.08 |
| HSPA8 | Y | Y | 1.07 |
| SLC3A2 | Y | Y | 1.07 |
| ENO1 | Y | Y | 1.05 |
| PRDX1 | Y | Y | 1.03 |
| GAPDH | Y | Y | 1.03 |
| ANXA11 | Y | Y | 1.02 |
| PDCD6IP | Y | Y | 1.01 |
| BSG | Y | Y | 1.00 |
| Uba1 | Y | Y | 0.99 |
| FLOT1 | Y | Y | 0.99 |
| ADAM10 | Y | Y | 0.99 |
| TLN1 | Y | Y | 0.96 |
| TPI1 | Y | Y | 0.96 |
| RALA | Y | Y | 0.95 |
| VCL | Y | Y | 0.95 |
| ACLY | Y | Y | 0.94 |
| ATP1A1 | Y | Y | 0.94 |
| PRDX2 | Y | Y | 0.94 |
| ALDOA | Y | Y | 0.94 |
| ACTN4 | Y | Y | 0.93 |
| ANXA1 | Y | Y | 0.93 |
| ACTN1 | Y | Y | 0.93 |
| ANXA6 | Y | Y | 0.92 |
| ANXA5 | Y | Y | 0.92 |
| RAB7A | Y | Y | 0.91 |
| TFRC | Y | Y | 0.90 |
| GPI | Y | Y | 0.90 |
| PGAM1 | Y | Y | 0.89 |
| ANXA7 | Y | Y | 0.87 |
| GDI2 | Y | Y | 0.87 |
| CAP1 | Y | Y | 0.87 |
| HSPA1A | Y | Y | 0.87 |
| FASN | Y | Y | 0.86 |
| YWHAG | Y | Y | 0.85 |
| TUBB4B | Y | Y | 0.84 |
| EZR | Y | Y | 0.84 |
| TSG101 | Y | Y | 0.84 |
| PKM | Y | Y | 0.83 |
| CDC42 | Y | Y | 0.83 |
| CFL1 | Y | Y | 0.82 |
| RAB5C | Y | Y | 0.80 |
| MSN | Y | Y | 0.80 |
| PGK1 | Y | Y | 0.79 |
| RAB10 | Y | Y | 0.79 |
| YWHAE | Y | Y | 0.78 |
| Rac1 | Y | Y | 0.77 |
| ITGB1 | Y | Y | 0.74 |
| YWHAB | Y | Y | 0.74 |
| RAP1B | Y | Y | 0.73 |
| YWHAZ | Y | Y | 0.70 |
| YWHAQ | Y | Y | 0.68 |
| PFN1 | Y | Y | 0.66 |
| RHOA | Y | Y | 0.61 |
| GSN | Y | Y | 0.61 |
| EHD1 | Y | Y | 0.60 |
| GNAS | Y | Y | 0.59 |
| ACTB | Y | Y | 0.59 |
| CD9 | Y | Y | 0.59 |
| GNB2 | Y | Y | 0.58 |
| PPIA | Y | Y | 0.57 |
| HLA-A | Y | Y | 0.57 |
| GNAI2 | Y | Y | 0.53 |
| GNB1 | Y | Y | 0.53 |
| CLIC1 | Y | Y | 0.50 |

**Supplementary Table 2: EV proteins that are significantly different in EV preparations from MDA-MB-231 cells cultured in either Opti-MEM or EV-DEP DMEM.**

| Species/protein ID | Adjusted p-value |
|---|---|
| Bovine Hemoglobin subunit alpha | 9.40098E-06 |
| Human Serglycin | 0.000556071 |
| Human UHRF1-binding protein 1 | 0.000721796 |
| Bovine Alpha-1-antiproteinase | 0.000755283 |
| Human Histone H1.2 | 0.001093171 |
| Human Histone H3.1; H3.3; H3.2; H3.1t; H3.3C | 0.001609695 |
| Human Thrombospondin-1 | 0.001609695 |
| Human Histone H2B type 1-N | 0.002365533 |
| Bovine Serpin A3-1 | 0.002901971 |
| Human Histone H1.0; H1.0, N-terminally processed | 0.004579547 |
| Human Histone H2AX | 0.004579547 |
| Bovine serpin A3-7-like precursor | 0.004579547 |
| Human Histone H2A type 2-B | 0.005810001 |
| Bovine Apolipoprotein E | 0.005810001 |
| Bovine Hemoglobin fetal subunit beta | 0.005810001 |
| Human Histone H2A.V; H2A.Z | 0.006026549 |
| Human Core histone macro-H2A.1 | 0.006712778 |
| Human Histone H2B type 1-C/E/F/G/I | 0.00673051 |
| Bovine Inter-alpha-trypsin inhibitor HC2 component homolog | 0.008201655 |
| Human Histone H2A type 1-D | 0.010820638 |
| Human Histone H4 | 0.010835622 |
| Human Thioredoxin-related transmembrane protein 1 | 0.01085701 |
| Human Thymidylate synthase | 0.012974637 |
| Bovine Clusterin | 0.012974637 |
| Bovine Vitronectin | 0.012974637 |
| Bovine Albumin | 0.01438022 |
| Human U1 small nuclear ribonucleoprotein 70 kDa | 0.017398326 |
| Bovine Fetuin-B | 0.017423475 |
| Bovine Prothrombin | 0.019550083 |
| Bovine immunoglobulin heavy constant mu | 0.023454327 |
| Human Kunitz-type protease inhibitor 2 | 0.024679111 |
| Human Protein CYR61 | 0.024679111 |
| Bovine Alpha-2-HS-glycoprotein | 0.024679111 |
| Human Histone H2B type 2-E | 0.025915919 |
| Human Transmembrane protein 109 | 0.028617093 |
| Human Transmembrane emp24 domain-containing protein 10 | 0.03006098 |
| Human Carbonic anhydrase 12 | 0.033540835 |
| Human Ubiquitin-fold modifier-conjugating enzyme 1 | 0.033540835 |
| Human Vesicle transport protein GOT1B | 0.033540835 |
| Human Serine protease HTRA1 | 0.040141311 |
